# Phytochemical characterization and *in vitro* antibacterial activity of *Xeroderris stuhlmannii* (Taub.) Mendonca & E.P. Sousa bark extracts

**DOI:** 10.1101/2020.10.29.360776

**Authors:** Major A. Selemani, Luckmore F. Kazingizi, Emily Manzombe, Lorraine Y. Bishi, Cleopas Mureya, Tichaziwa T. Gwata, Freeborn Rwere

**Affiliations:** Department of Chemistry, School of Natural Sciences and Mathematics, Chinhoyi University of Technology, Chinhoyi, Zimbabwe; Tobacco Research Board, Kutsaga Research Station, Airport Ring Road, Harare, Zimbabwe

**Keywords:** *Xeroderris stuhlmannii* (Taub.) Mendonca & E.P. Sousa, phytochemicals, LC-MS/MS, GC-MS, Minimum Inhibitory Concentrations (MICs)

## Abstract

The use of herbal medicine is common in many rural communities in Zimbabwe because generic drugs are expensive and not readily available. In this work, we documented the important phytochemicals from *Xeroderris stuhlmannii* (Taub.) Mendonca & E.P. Sousa (Murumanyama in Shona), bark extracts and tested their antibacterial activity in order to demonstrate its potential as an antimicrobial agent. Qualitative screening of secondary metabolites confirmed the presence of alkaloids, flavonoids, terpenoids, steroids and polyphenols in the crude bark extracts. The MICs values for the crude extracts on six bacterial strains ranged from 0.23–0.80 mg/mL. Antimicrobial tests showed higher potency for crude bark extracts on *E. Coli* (MIC, 0.232 mg/mL) and lower potency on *coliform* (MIC, 0.798 mg/mL). LC-MS/MS analysis of various fractions confirmed the presence of twenty-eight phytochemicals whereas, twelve phytochemicals were identified using GC-MS. Both techniques confirmed the presence of ursolic acid, roburic acid, reticuline, rotenone and p-coumaric acid glucoside in hexane and methanol extracts. In summary, our findings show that *Xeroderris stuhlmannii* (Taub.) Mendonca & E.P. Sousa contain many phytochemical compounds that have antimicrobial activity. Moreover, some of the compounds in the bark extract have been shown to possess antioxidant, antiviral, antitumor and anti-inflammatory properties. Thus, *Xeroderris stuhlmannii* (Taub.) Mendonca & E.P. Sousa barks can act as a useful herbal supplement for treatment of a number of diseases in rural communities where modern drugs are expensive and not readily available.

## 1. INTRODUCTION

Over the years, focus on plant research has increased all over the world and a large body of evidence has been collected to show the immense potential of medicinal plants used in traditional systems in African and beyond (Barbieri et al., 2017; Cargnin and Gnoatto, 2017; Cock et al., 2019; Haudecoeur et al., 2018; Maroyi, 2013; Njerua et al., 2013). Scientific interest in medicinal plants has also been necessitated by the increased efficiency of new plant derived drugs and rising concerns about the side effects associated with the modern synthetic drugs. Herbal medicine offers the advantage that it is cheap, safe and readily available in rural communities. Medicinal plants are a rich source of bioactive phytochemicals and/or bio-nutrients (Mugumbate et al., 2018). These bioactive phytochemicals possess antibacterial, antiviral, antitumor, antidiabetic, anti-inflammatory, anticancer, antimalarial, antioxidant, antimigraine and antiparasitic properties (Barbieri et al., 2017; Cargnin and Gnoatto, 2017; Cock et al., 2019; Haudecoeur et al., 2018).

*Xeroderris stuhlmannii* (Taub.) Mendonca & E.P. Sousa (leguminosae family) has long been used as herbal medicine to treat a number of ailments such as coughs, diarrhea, malaria, colds, rheumatoid arthritis, stomachache, dysentery and eye infections (Maroyi, 2013; Ngarivhume et al., 2015). It is well known for its diverse biological and pharmacological properties. A survey by Ngarivhume et. al., in rural communities in Zimbabwe has shown that the roots and barks of *Xeroderris stuhlmannii* (Taub.) Mendonca & E.P. Sousa are used to treat colds and stomach pains (Ngarivhume et al., 2015). Studies by Asase et al., have shown the antimalarial potential of *Xeroderris stuhlmannii* (Taub.) Mendonca & E.P. Sousa (Asase et al., 2005). However, there are no systematic studies that have been carried out to date on *Xeroderris stuhlmannii* (Taub.) Mendonca & E.P. Sousa in order to identify its bioactive phytochemicals that are responsible for these properties.

The availability of various modern analytical techniques has facilitated the extraction, isolation and identification of the biologically active phytochemicals from plant extracts. Liquid chromatography (LC) and gas chromatography (GC) coupled with mass spectroscopy (MS), (LC-MS, LC-MS/MS and GC-MS) and nuclear magnetic resonance (NMR) spectroscopy are important spectroscopic techniques that give detailed structural information about the nature of different phytochemicals in plant extracts. Combination of these analytical techniques (LC/MS or GC/MS) has successfully been used to isolate the bioactive compounds from plants and establish their chemical structures (Faraone et al., 2018; Fu et al., 2014; Keskes et al., 2017). However, precise identification of all compounds in crude plant extracts can be a complex task as they contain a wide variety of structures. In this work, we have employed liquid chromatography tandem mass spectrometry (LC-MS/MS) and gas chromatography mass spectroscopy (GC-MS) to identify the phytochemicals from *Xeroderris stuhlmannii* (Taub.) Mendonca & E.P. Sousa because these techniques have been proven to be efficient tools for rapid identification and quantification of known and unknown compounds in complex systems (Fu et al., 2014).

Based on indigenous knowledge, we hypothesized that *Xeroderris stuhlmannii* (Taub.) Mendonca & E.P. Sousa contains a plethora of phytochemicals that are responsible for treatment of bacterial infections. Thus, current research was aimed at identifying bioactive phytochemical compounds in bark extracts of *Xeroderris stuhlmannii* (Taub.) Mendonca & E.P. Sousa and to test their antimicrobial activity. The results of this work revealed the presence of a number of important alkaloids, flavonoids, tepernoids and tri-tepernoids, polyphenols, steroids and other phytochemicals. Antibacterial tests on both the crude and purified extracts have shown the potential of bark extracts to fight a number of Gram-negative and Gram-positive bacteria.

## 2. MATERIALS AND METHODS

### 2.1 Materials

The *Xeroderris stuhlmannii* (Taub.) Mendonca & E.P. Sousa (murumanyama in Shona) barks were obtained from Domboshava, Mashonaland East Province near Harare, Zimbabwe after getting permission from the local religious headmen and in accordance with Zimbabwe laws governing the use of traditional plants for research. The samples were transported to Kutsaga Research Station for drying, storage and laboratory analysis. Analytical grade reagents such as methanol, acetone, ethyl acetate, diethyl ether, hexane, chloroform, sulphuric acid, ferric chloride and hydrochloric acid were purchased from Sigma-Aldrich and Merck Chemicals (Pty) Ltd, South Africa via our local vendors.

### 2.2 Methods

#### 2.2.1 *Extraction of phytochemicals from Xeroderris stuhlmannii* (Taub.) Mendonca & E.P. Sousa *bark samples*

Plant samples (bark) were pre-washed first with running water followed by distilled water. The samples were dried at ambient room temperature. Once the plant samples were fully dried, they were grinded into fine powder using an electric grinder. Sieving was then done through a 1 mm sieve to obtain finely divided powdered samples. The powder samples were stored in a cool dark place in opaque bottles before analysis. Crude bark extracts were obtained using the maceration method by soaking 5 g of the powdered plant material in 50 mL of different solvents; water, methanol, acetone, diethyl ether, ethyl acetate and hexane, in sterile screw capped glass containers. After 24 h, the samples were macerated for ~30-45 min using a macerator and then filtered using the Whatman filter paper (No. 541) under gravity to collect the extracts. The extracts were collected in opaque containers and stored at 4°C in the cold room before analysis. The average yield of crude extracts was between 13-15%.

#### 2.2.2 *Qualitative phytochemical screening of Xeroderris stuhlmannii* (Taub.) Mendonca & E.P. Sousa *bark extracts*

Qualitative phytochemical screening of secondary metabolites was carried out to test for the presence of alkaloids, flavonoids, di-terpenoids and tri-terpenoids, saponins, phenolics and polyphenolics, coumarins, quinones, glycosides, steroids, amino acids, proteins, tannins and carbohydrates using the published procedure by (Riddhu. and Payal., 2015).

#### 2.2.3 *Antimicrobial activity tests for Xeroderris stuhlmannii* (Taub.) Mendonca & E.P. Sousa *bark extracts*

Antibacterial activities of the extracts against six bacterial strains (*S. aureus, P. aeruginosa, E. coli, Coliform, Listeria spp* and *Salmonella spp*) were evaluated using disk diffusion method on Mueller-Hinton agar (MHA) and the zones of inhibition are reported in millimeter (mm). Specifically, three (3) to five (5) well isolated colonies from an agar plate were placed in 5 mL of tryptic soy broth and incubated at 35 °C until the turbidity of the growth was comparable to the 0.5 McFarland standards (usually 2-6 h). The Mueller-Hinton agar plates were inoculated with bacterial strains under sterile conditions. Sterile disks (diameter = 2 mm) were filled with ~5 μL (~0.35 mg/disc) of the plant extracts and placed on top of the plates. The plates were incubated at 37°C for ~24 h. After incubating for 24 h, the diameters of the growth inhibition zones were measured using a digital Vernier caliper. All tests were performed in triplicates and on two (2) separate days in order to validate the results.

#### 2.2.4 *Determination of Minimum Inhibitory Concentrations (MIC’s) for Xeroderris stuhlmannii* (Taub.) Mendonca & E.P. Sousa *bark extract*

Approximately, 700 mg of dry plant extract was dissolved in 10 mL of methanol to make up a concentration of ~70 mg/mL which was then taken as the stock solution for serial dilutions. Different concentrations of the bark extract (70, 35, 17.5, 8.75, and 4.375 mg/mL) were prepared from the stock solution. Mueller-Hilton agar was poured into sterile Petri dishes and seeded with bacterial suspensions of the pathogenic strains. The loaded filter paper discs (2 mm) with different concentrations of the effective plant extract were placed on the top of the Mueller-Hinton agar plates. The plates were incubated at 37 °C for 24 h. After 24 h, the inhibition zones were measured using a digital Vernier caliper and recorded against the concentrations. The MIC’s were determined using the linear diffusion and quadratic models according to the published procedures by Parisot and coworkers (Bonev et al., 2008).

#### 2.2.5 Purification, identification and characterization of phytochemicals using liquid chromatography tandem mass spectrometry (LC-MS/MS)

The methanolic bark extracts were obtained using the maceration process as described in section 2.2.1. The methanol extracts were purified on a Gel permeation chromatography (GPC) instrument by collecting different fractions at time different intervals. The GPC stationary phase was prepared from s-x3 Bio-beads. Phytochemical constituents in purified plant extracts were determined using a state-of-the art liquid chromatography tandem spectroscopy (LC-MS/MS) in electrospray ionization mode as described below. An AB Sciex 5500 model tandem mass spectrometer (MS/MS) with 4 software’s (i.e. Analyst 1.6.2, Peak View 2.2, Library View 1.0.1 and Master View 1.1) coupled to Agilent Liquid Chromatography (LC) 1260 series system was used to analyze the purified fractions. The Agilent LC system was made up of the **v**acuum degasser GG1379 B, binary pump G1312B, auto-sampler G1367C, column oven G1316B, analytical column (phenomenex synergi 4 μ Fusion-RP 100 Å, 50 x 2 mm) and guard column (phenomenex security guard cartridge with Fusion-RP 4 x 2.00 mm cartridge). Mass spectrometry was performed using electrospray ionization. The separation of phytochemicals was accomplished on a reverse-phase (50 x 2 mm) C18 column with a mobile phase of methanol/water (90/10%) and 5 mM ammonium formate.

#### 2.2.6 Identification and characterization of phytochemicals using gas chromatography mass spectrometry (GC-MS)

Crude bark extracts in hexane were obtained by the maceration process. The extracts were chromatographed on a silica gel column in order to obtain different fractions that were analyzed by GC-MS. GC-MS analysis of volatile phytochemical compounds present in the bark extracts of the plant was carried out using an Agilent GC MS model number 5977B comprising of an AOC-20i auto sampler and gas chromatography interfaced to mass spectrometer in the following conditions: Column Elite-1 fused silica capillary (30×0.25 mm x 1D x 1μm of capillary column composed with 100% siloxane) operating in electron ionization system with an impart mode of 70 eV, helium (99.999 %) as carrier gas, flow rate of 1 mL/min, injector temperature 250°C, oven temperature from 40°C (Isothermal for 5 min with an increase of 10°C to 300°C/min). Running time was 24 min.

### 2.3 Statistical Analysis

Statistical analysis was performed using GraphPad Prism 8.0 by ANOVA and Tukey’s post hoc with significance set at P<0.05. All data are presented as mean ± s.d.

## 3. RESULTS

### 3.1 *Yield of Xeroderris stuhlmannii* (Taub.) Mendonca & E.P. Sousa *bark extracts*

The average percentage yield of crude bark extract was ~14%. The very low percentage yield of the extract is consistent with the literature that showed low yield for phytochemicals in most of the studied plants (Sultana et al., 2009). Sultana et al., showed that even though the extract yields of the plant materials are strongly dependent on the nature of extracting solvent and method used, their overall percentage yields are generally low (Sultana et al., 2009).

### 3.2 *Qualitative Analysis of plant extracts of Xeroderris stuhlmannii* (Taub.) Mendonca & E.P. Sousa

Phytochemical screening of *Xeroderris stuhlmannii* (Taub.) Mendonca & E.P. Sousa bark extracts was carried out in various solvents to determine the presence or absence of secondary metabolites. As shown in Table 1, qualitative screening of secondary metabolites in *Xeroderris stuhlmannii* (Taub.) Mendonca & E.P. Sousa bark extracts revealed the presence of tannins, saponins, terpenoids, quinones, coumarins, glycosides, reducing sugars, carbohydrates, alkaloids, flavonoids, phenols, proteins and amino acids. Qualitative phytochemical studies in Table 1 also showed that methanol extracted the widest range of phytochemicals compared to other solvents. Methanol extracts had high levels of alkaloids, phenolics, terpenoids and flavonoids in abundance. Diethyl ether had high levels of terpenoids, coumarins and phenolics. Ethyl acetate and water had moderate levels of secondary metabolites, whereas hexane and acetone extracts had the least number of secondary metabolites in abundance.

**Table1.** Qualitative Screening of the Secondary Metabolites in *Xeroderris stuhlmannii* (Taub.) Mendonca & E.P. Sousa *Bark* Extracts

| Secondary metabolite | Water | Methanol | Acetone | Ethyl Acetate | Diethyl Ether | Hexane |
| --- | --- | --- | --- | --- | --- | --- |
| Alkaloids | + | ++ | - | + | + | + |
| Flavonoids | - | ++ | + | - | - | + |
| Saponins | ++ | + | - | - | - | - |
| Tannins | ++ | + | + | - | + | - |
| Terpenoids | - | ++ | - | + | ++ | + |
| Phenols | + | ++ | - | - | ++ | - |
| Coumarins | - | - | + | - | ++ | - |
| Quinones | - | + | + | - | - | - |
| Glycosides | - | + | - | - | - | - |
| Carbohydrates | + | + | + | + | + | + |
| Steroids | + | + | - | - | - | + |
| Reducing sugars | + | + | - | ++ | - | + |
| Proteins and Amino acids | - | ++ | + | + | + | - |

**KEY:** - not detectable + trace amount ++ abundant

### 3.3 *Antimicrobial activity of the Xeroderris stuhlmannii* (Taub.) Mendonca & E.P. Sousa *bark extracts*

*In vitro* antimicrobial activity of bark extracts was determined using the agar disc diffusion method, an assay that has long been used as a qualitative technique for analyzing the antimicrobial activity of plant extracts because of its simplicity and rapidity. The methanolic extracts were evaluated against six (6) food poisoning Gram-negative and Gram-positive bacterial strains namely; *Staphylococcus aureus, Pseudomonas aeruginosa, Escherichia Coli, Coliform, Listeria* and *Salmonella* and the results are summarized in Table 2. Antimicrobial results in Table 2 demonstrate that the extracts were capable of inhibiting the bacterial strains with zones of inhibition ranging from 4.91 to 6.5 mm at highest concentration of extract. The *Salmonella* strain (6.5 mm) was most inhibited, whereas the *coliform* strain (4.91 mm) was least inhibited i.e., most resistant strain (Table 2). The results show that *coliform* was the most resistant strain to the bark extracts followed by *E. coli* whereas, *S. aureus, Salmonella spp* and *P. aeruginosa* were the most susceptible strains to the plant extracts.

**Table 2.** Zones of inhibition (in mm) of six clinical bacterial strains at different concentrations of crude bark extract

| Conc<br>(mg/mL) | Bacterial Strain |  |  |  |  |  |
| --- | --- | --- | --- | --- | --- | --- |
|  | *Salmonella*<br>*spp* | *Listeria spp* | *Coliform* | *Escherichia*<br>*Coli* | *Pseudomonas*<br>*Aeruginosa* | *Staphylococcus.*<br>*Aureus* |
| 70.0 | 6.50 ± 0.03 | 5.96 ± 0.20 | 4.91 ± 0.01 | 5.58 ± 0.08 | 6.06 ± 0.08 | 6.33 ± 0.04 |
| 35.0 | 6.05 ± 0.10 | 5.70 ± 0.03 | 4.06 ± 0.08 | 4.84 ± 0.01 | 5.05 ± 0.23 | 5.68 ± 0.04 |
| 17.5 | 4.94 ± 0.02 | 4.48 ± 0.04 | 3.24 ± 0.01 | 4.10 ± 0.16 | 4.53 ± 0.06 | 4.53 ± 0.01 |
| 8.9 | 3.89 ± 0.03 | 3.16 ± 0.08 | 2.68 ± 0.04 | 3.48 ± 0.04 | 3.75±0.10 | 3.11 ± 0.15 |
| 4.4 | 3.04 ± 0.08 | 2.65 ± 0.04 | 0 | 2.91 ± 0.06 | 2.70±0.15 | 2.88 ± 0.04 |

Minimum inhibitory concentration (MIC) values of the bark plant extracts were determined using the disc diffusion method to evaluate the effectiveness of the plant extracts against the six bacterial strains (Table 3). The data was fitted using the linear (Eq. 1) and quadratic (Eq. 2) diffusion models according to work by Boven and Parisot (Bonev et al., 2008). The MICs were estimated from plots of *x* or *x*^2^ versus ln(c) and are summarized in Table 3. As shown in Table, 3, the bark extracts had antibacterial activity against both Gram-positive and Gram-negative bacteria, with MIC values ranging from 0.23–0.80 mg/mL. The methanolic bark extract had the greatest activity against *E. coli* (0.232 mg/mL), *P. aeruginosa* 0.384 (mg/mL) and *Salmonella spp* (0.421 mg/mL). The least activity (0.632-0.798 mg/L) was observed for *S. aureus, coliform and Listeria spp.* Small MICs presented here suggested high bacterial efficacy. The high efficacy of plant extracts indicates the presence of antibacterial phytochemicals in *Xeroderris stuhlmannii* (Taub.) Mendonca & E.P. Sousa.

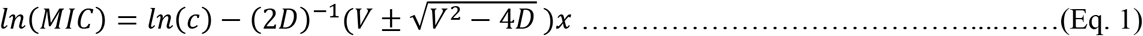

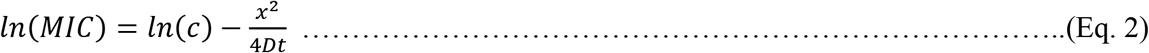

where *D* is the diffusion coefficient, c is concentration (mg/mL), *x* is zone of inhibition (mm), *V* is a coefficient characterizing the dissipation rate and t is the time of crude extract diffusion.

**Table 3.** Minimum inhibitory concentrations (MIC’s) of the bark *Xeroderris stuhlmannii* (Taub.) Mendonca & E.P. Sousa extract against Gram-negative and Gram-positive bacteria

| Parameter | Bacterial Strain |  |  |  |  |  |
| --- | --- | --- | --- | --- | --- | --- |
|  | *Salmonella*<br>*app* | *Listeria*<br>*spp* | *Coliform* | *E. Coli* | *P.*<br>*aeruginosa* | *S. aureus* |
| *x/ln(c) model |  |  |  |  |  |  |
| MIC (mg/mL) | 0.421 | 0.632 | 0.798 | 0.232 | 0.384 | 0.648 |
| (R<sup>2</sup>) | 0.986 | 0.961 | 0.992 | 0.997 | 0.989 | 0.965 |
| x<sup>2</sup>/ln(c) model |  |  |  |  |  |  |
| MIC (mg/mL) | 2.32 | 2.80 | 4.18 | 1.85 | 2.32 | 2.94 |
| (R<sup>2</sup>) | 0.985 | 0.957 | 0.972 | 0.978 | 0.981 | 0.959 |

\*x is zone of inhibition and c is concentration

### 3.4 LC-MS/MS Analysis of Bark Extracts

The retention times, names and mass spectral data for methanolic extracts of *Xeroderris stuhlmannii* (Taub.) Mendonca & E.P. Sousa bark area summarized in Table 4. As shown in Table 4, a total of twenty-eight (28) putative/tentative phytochemical compounds were identified based on matching the retention times and molecular fragmentation information from the LC-MS/MS experiments with Analyst, Library view and Master view libraries. The secondary metabolites in bark extracts were mainly alkaloids (4), flavonoids (7), terpenoids (7), phenolics (1), steroids (3), lipid (1) and glycoside (4). One compound was not classified.

**Table 4.** LC MS/MS analysis of *Xeroderris stuhlmannii* (Taub.) Mendonca & E.P. Sousa extracts

| Putative compound name | Formula | Parent mass (m/z) | Retention Time | Library Score (%) | Phytochemical classification |
| --- | --- | --- | --- | --- | --- |
| Octanoylphenylalaninate | $C_{17}H_{24}NO_3$ | 291.1830 | 7.17 | 95.0 | Alkaloid |
| Apiin | $C_{26}H_{28}O_{14}$ | 565.1543 | 8.63 | 85.0 | Flavonoid |
| Syringin | $C_{17}H_{24}O_9$ | 373.1493 | 9.55 | 90.0 | glycoside |
| Cotinine | $C_{10}H_{12}N_2O$ | 177.1018 | 10.1 | 76.0 | Alkaloid |
| Myricitrin | $C_{21}H_{20}O_{12}$ | 465.2021 | 10.94 | 89.4 | Flavanoid |
| 2'-Deoxyuridine-5'-triphosphate | $C_9H_{11}N_2O_{14}P_3$ | 464.9475 | 11.02 | 89.0 | Glycoside |
| Coriolic acid | $C_{18}H_{32}O_3$ | 297.2424 | 11.41 | 91.0 | Lipid |
| Caffeine | $C_8H_{10}N_4O_2$ | 195.0877 | 11.73 | 98.0 | Alkaloid |
| maackiain | $C_{16}H_{12}O_5$ | 285.0763 | 12.22 | 82.0 | Flavonoid |
| p-coumaric acid glucoside | $C_{15}H_{18}O_8$ | 327.1074 | 12.92 | 97.0 | Phenolic |
| Myricetin | C<sub>15</sub>H<sub>10</sub>O<sub>8</sub> | 319.1773 | 13.07 | 85 | Flavonoid |
| Gibberellin A1(-1) | C<sub>19</sub>H<sub>23</sub>O<sub>6</sub> | 348.1113 | 13.17 | 78 | Diterpenoid |
| (-)-sativan | C<sub>17</sub>H<sub>18</sub>O<sub>4</sub> | 287.1278 | 13.58 | 100 | Flavanoid |
| P1-uridyl-P2-phenyl<br>diphosphate | C<sub>15</sub>H<sub>16</sub>N<sub>2</sub>O<sub>12</sub>P<sub>2</sub> | 478.9305 | 13.66 | 93.8 | - |
| Retinoic acid | C<sub>20</sub>H<sub>28</sub>O<sub>2</sub> | 301.0785 | 14.39 | 99.5 | Tetraterpenoid |
| Coniferyl alcohol | C<sub>10</sub>H<sub>12</sub>O<sub>3</sub> | 181.0860 | 14.57 | 96.0 | Glucoside |
| (R or S)-reticuline | C<sub>19</sub>H<sub>24</sub>N<sub>1</sub>O<sub>4</sub> | 331.1786 | 14.92 | 75.0 | Alkaloid |
| Kaempferide | C<sub>16</sub>H<sub>12</sub>O<sub>6</sub> | 301.0707 | 15.04 | 100 | Flavonoid |
| 5-Amino-6-ribitylamino<br>uracil | C<sub>9</sub>H<sub>16</sub>N<sub>4</sub>O<sub>6</sub> | 277.2159 | 16.04 | 74.2 | Glycoside |
| castasterone | C<sub>28</sub>H<sub>48</sub>O<sub>5</sub> | 465.1987 | 16.93 | 89.4 | steroid |
| Gibberellin A12<br>aldehyde | C<sub>20</sub>H<sub>27</sub>O<sub>3</sub> | 316.1994 | 17.37 | 73.7 | Terpenoid |
| Khasianine | C<sub>39</sub>H<sub>63</sub>NO<sub>11</sub> | 722.4880 | 18.60 | 98.8 | Steroidal<br>alkaloid |
| Roburic acid | C<sub>30</sub>H<sub>48</sub>O<sub>2</sub> | 441.3779 | 18.94 | 81.7 | Triterpenoid |
| Ursolic acid | C<sub>30</sub>H<sub>48</sub>O<sub>3</sub> | 457.5084 | 18.95 | 92.0 | Triterpenoid |
| Oleanolic aldehyde | C<sub>30</sub>H<sub>48</sub>O<sub>2</sub> | 441.3702 | 19.52 | 81.7 | Triterpenoid |
| Garajonone | C<sub>22</sub>H<sub>32</sub>O<sub>9</sub> | 441.2002 | 19.05 | 81.7 | Diterpenoid |
| Brassinolide | C<sub>28</sub>H<sub>48</sub>O<sub>6</sub> | 481.3118 | 21.0 | 73.8 | steroid |
| Rotenone | C<sub>23</sub>H<sub>22</sub>O<sub>6</sub> | 395.1621 | 20.25 | 97.1 | Flavonoid |

Figure 1 summarizes the fragmentation pattern of selected plant metabolites. Based on fragmentation behaviors and literature we tentatively assigned several fragments to their respective ions. Ursolic acid exhibited major peaks at m/z 392.85, 410.8, 438.69 and 457.92 corresponding to C_29_H_45_^+^ C_29_H_4_7O^+^ C_30_H_47_O_2_^+^ and C_30_H_49_O_3_^+^ ions, respectively (Novotny et al., 2003). Roburic acid has peaks at m/z 370.98, 422,82 and 441.28 for C_27_H_46_^+^, C_30_H_47_O_2_^+^ and C_30_H_49_O_2_^+^ respectively. The flavonoid kaempferide has a molecular ion peak (M+H)^+^ at m/z 301.07 for the C_16_H_13_O_6_^+^ ion. Other major peaks for kaempferide occur at m/z of 269.56 and 282.37 and likely correspond to C_15_H_10_O_5_^+^ and C_16_H_11_O_5_^+^ ions, respectively (Aghakhani et al., 2018). Other identified compounds were the broad-spectrum insecticide rotenone with a molecular ion peak at m/z 395.29 for the C_23_H_23_O_6_^+^ ion. Other major peaks for rotenone occur at m/z 352.83, 358.47 and 376.59. The peak at m/z 192 of elemental composition C_11_H_12_O_3_^+^., for rotenone was previously assigned to the 6,7-dimethoxy–2H-1-benzo-pyran radical cation (Chen et al., 2015).

**Figure 1.**
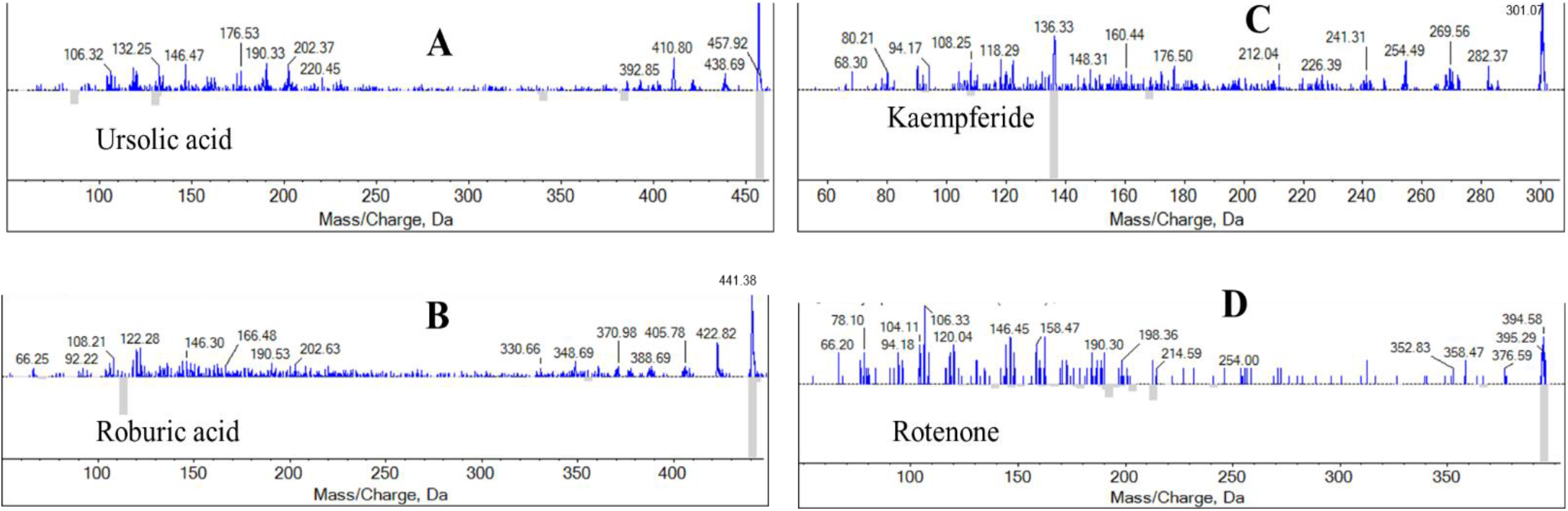
The MS/MS spectra of selected phytochemicals with antibacterial, anti-inflammatory, anti-tumor and anti-parasitic properties.

### 3.5 GC MS analysis of bark extracts

The GC MS chromatogram of n-hexane bark extract (Figure 2, Trace A) showed nine (9) major well-resolved peaks and one broad peak near 5 min. The broad peak near 5 min was further chromatographed on silica gel column and five (5) additional compounds were analysis using GC MS (Figure 2, Traces B-F). The major compounds in hexane extract were identified based on matching the retention times, molecular weight and NIST Library from GC-MS and are summarized in Table 5. The secondary metabolites in the extract were mostly alkaloids (7), terpenoids (3), flavonoids (2), phenol (1) and quinolines (1). Some of the compounds identified by GC MS were also present in methanolic extract that was studied using LC MS/MS. The common compounds are ursolic acid, roburic acid, apiin, rotenone and reticuline. Structures of some of these common phytochemicals and additional compounds are shown in Figure 3.

**Figure 2.**
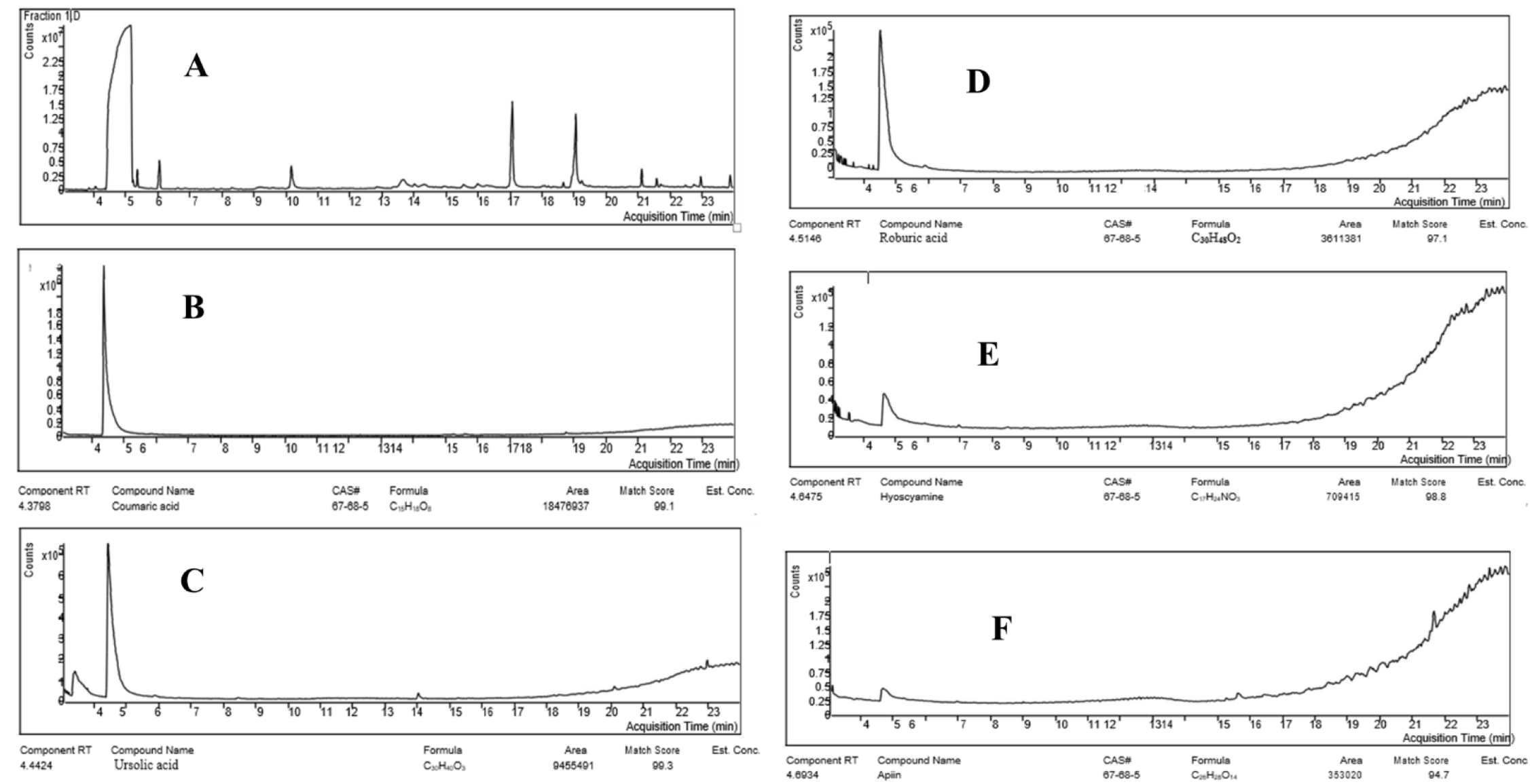
Chromatographic profiles of the fractions of hexane extracts of *Xeroderris stuhlmannii* (Taub.) Mendonca & E.P. Sousa. Fraction with all compounds, A; p-coumaric acid glucoside, B; ursolic acid, C; roburic acid, D; hysocyamine, D; and apiin, F.

**Figure 3.**
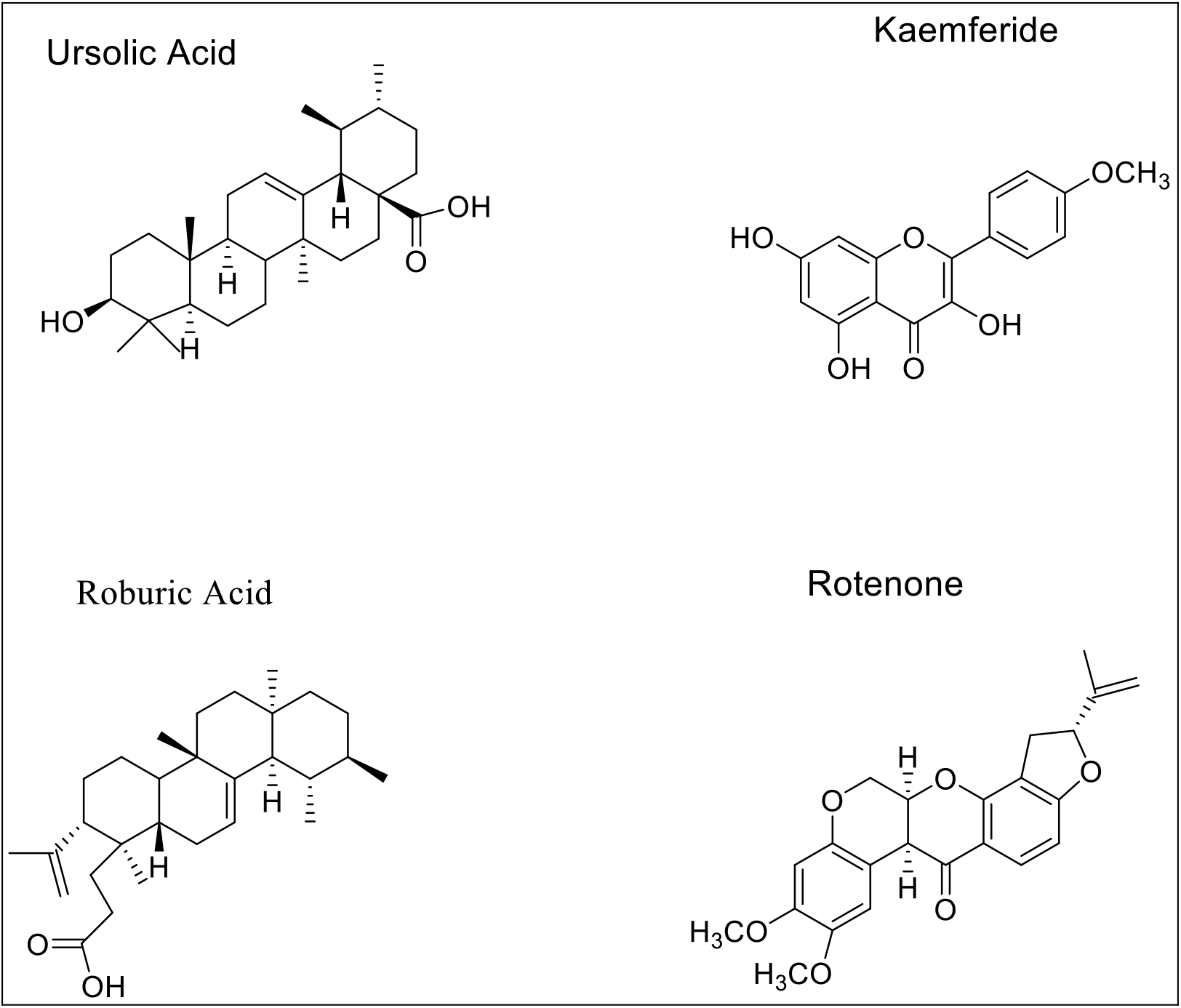
Structures of selected phytochemicals with antibacterial, anti-inflammatory, anti-tumor and anti-parasitic properties.

**Table 5.** GC MS analysis of *Xeroderris stuhlmannii* (Taub.) Mendonca & E.P. Sousa

| Putative compound name | Formula | Parent mass (m/z) | Retention time | Library score (%) | Phytochemical classification |
| --- | --- | --- | --- | --- | --- |
| p-coumaric acid glucoside | C<sub>15</sub>H<sub>18</sub>O<sub>8</sub> | 327.1074 | 4.38 | 99.1 | phenolic |
| Ursolic acid | C<sub>30</sub>H<sub>48</sub>O<sub>3</sub> | 457.5084 | 4.44 | 99.3 | Triterpernoid |
| Roburic acid | C<sub>30</sub>H<sub>48</sub>O<sub>2</sub> | 441.3779 | 4.51 | 97.1 | Triterpernoid |
| Hyoscyamine | C<sub>17</sub>H<sub>24</sub>NO<sub>3</sub> | 291.1830 | 4.65 | 98.8 | Alkaloid |
| Apiin | C<sub>26</sub>H<sub>28</sub>O<sub>14</sub> | 565.1543 | 4.69 | 94.7 | Flavonoid |
| Rotenone | C<sub>23</sub>H<sub>22</sub>O<sub>6</sub> | 395.162 | 5.01 | 97.01 | Flavonoid |
| Robustaquinone E | C<sub>16</sub>H<sub>11</sub>O<sub>7</sub> | 316.199 | 5.15 | 75.0 | Quinoline |
| (S)-Coclaurine | C<sub>17</sub>H<sub>20</sub>NO<sub>3</sub> | 287.152 | 6.01 | 86.0 | Alkaloid |
| (R)-Coclaurine | C<sub>17</sub>H<sub>20</sub>NO<sub>3</sub> | 287.152 | 10.30 | 86.0 | Alkaloid |
| (S)-reticuline | C<sub>19</sub>H<sub>24</sub>N<sub>1</sub>O<sub>4</sub> | 331.178 | 17.08 | 75.0 | Alkaloid |
| (R)-reticuline | C<sub>19</sub>H<sub>24</sub>N<sub>1</sub>O<sub>4</sub> | 331.178 | 18.85 | 75.0 | Alkaloid |
| Vindolinium | C<sub>25</sub>H<sub>33</sub>N<sub>2</sub>O<sub>6</sub> | 458.241 | 19.07 | 82.0 | Alkaloid |
| Geranylgeranyl diphosphate | C<sub>20</sub>H<sub>36</sub>O<sub>7</sub>P<sub>2</sub> | 451.200 | 21.24 | 76.0 | Terpenoid |
| Codeine | C<sub>18</sub>H<sub>22</sub>NO<sub>3</sub> | 301.167 | 23.01 | 99.5 | Alkaloid |

## 4. DISCUSSION

The use of plant-based traditional medicine in Africa remains an integral part of rural life because medicinal plants have many important phytochemicals and are easily accessible in many communities. Modern medicine is not readily accessible and affordable in those communities. In this work, we studied a medicinal plant called *Xeroderris stuhlmannii* (Taub.) Mendonca & E.P. Sousa because it has extensively been used to treat a number of ailments in Zimbabwe (Ngarivhume et al., 2015). Qualitative screening of *Xeroderris stuhlmannii* (Taub.) Mendonca & E.P. Sousa bark extracts showed that it contains several secondary plant metabolites such as alkaloids, flavonoids and terpenoids. We also carried out antimicrobial tests on six selected bacterial strains and showed that the crude bark extracts of *Xeroderris stuhlmannii* (Taub.) Mendonca & E.P. Sousa were very effective against these strains with MICs ranging from 0.23– 0.80 mg/mL. The antibacterial activity of crude extracts is likely due to the presence of one compound or a number of compounds that act synergistically. We used the theoretical framework model for analysis of agar diffusion assays data in order to determine the MICs for methanolic bark extracts. In this model, the zones of inhibition were fitted to the linear and quadratic diffusion models according to equations illustrated in the results section (Bonev et al., 2008). The linear diffusion model produced the best fit with the highest R^2^ values and low MICs. The major limitation to these two models is that the MICs values for two models differ from each other by ~5-fold. However, the linear model remains the best model because it predicts a linear dependence of indicator zone sizes on the logarithm of concentration of the diffusing agent and improves the accuracy of the results compared to the quadratic method (Bonev et al., 2008).

We also carried out LC MS/MS and GC MS analysis of chromatographed fractions of methanol and hexane extracts in order to identify the important phytochemicals. Our results confirmed the presence of several phytochemicals in *Xeroderris* stuhlmannii (Taub.) Mendonca & E.P. Sousa bark extracts. Identification of compounds was made possible by carefully analyzing the fragmentation patterns and comparison with published literature (Aghakhani et al., 2018; Chithalen et al., 2002; Novotny et al., 2003). Most notably, our LC/MS/MS and GC/MS results showed that ursolic acid, roburic acid, apiin, rotenone and reticuline were present in hexane and methanol extracts (See Table 4 and 5). The major classes of compounds identified in these studies were alkaloids, flavonoids and terpenoids. Many compounds in these different classes have well documented antibacterial properties that are presented below.

Alkaloids are organic nitrogenous bases with diverse and important physiological effects on humans and other animals (Barbieri et al., 2017). Alkaloids possess antibacterial, antimalarial, antiasthma, antihypertensive, antitumor and antiarrhythmic effects. At least ten (10) alkaloids were found in hexane and methanol extracts of *Xeroderris* stuhlmannii (Taub.) Mendonca & E.P. Sousa. Of these alkaloids is caffeine, a central nervous stimulant with some antibacterial properties (Sledz et al., 2015). GC/MS confirmed the presence of hyoscyamine, an alkaloid that acts as an antagonist of muscarinic acetylcholine receptors and has particularly been useful for control of neuropathic and chronic pain. Codeine is a narcotic pain-reliever and that has long been used as cough suppressant. Cotinine (m/z=177.1018) is an alkaloid that has been shown to prevent working and reference memory loss in a mouse model of Alzheimer’s Disease (Grizzell and Echeverria, 2014). The alkaloid, reticuline possesses potent central nervous system depressing effects (Morais et al., 1998).

Terpenoids are a large class of compounds found in plants with antibacterial, anti-inflammatory and antitumor activities (Barbieri et al., 2017). Approximately, eight di- and triterpenoids were found in *Xeroderris stuhlmannii* (Taub.) Mendonca & E.P. Sousa crude extracts. Ursolic acid is a triterpenoid with functional properties such as antibacterial, antiprotozoal and anti-inflammatory (Cargnin and Gnoatto, 2017). Other research has shown that ursolic acid has anti-obesity properties and can be a potential drug for therapeutic treatment of obesity. Ursolic acid also possesses anti-proliferative activity and can be used to treat cancer (Cargnin and Gnoatto, 2017; Navin and Mi Kim, 2016). LC MS/MS and GC MS of methanol and hexane extracts confirmed the presence of roburic acid at m/z, 441.3779. Roburic acid is a tetracyclic triterpenoid that possesses anti-inflammatory properties. Oleanolic aldehyde, a pentacyclic triterpenoid and hydroxy-aldehyde is widely found in plants and can be used for the treatment and management of chronic diseases. Studies by Karygianni et al. showed that oleanolic aldehyde possesses some antibacterial properties (Karygianni et al., 2019).

LC MS/MS and GC MS also confirmed the presence of eight important flavonoids in methanolic and hexane extracts of *Xeroderris stuhlmannii* (Taub.) Mendonca & E.P. Sousa (Tables 4 and 5). Kaempferide (m/z=301.07) is an O-methylated flavonoid with a broad range of properties including antibacterial, anticarcinogenic, anti-inflammatory, antioxidant, and antiviral properties (Eumkeb et al., 2012). Kaempferide has also been shown to inhibit pancreatic cancer growth by blockading an EGFR-related pathway. Padayachee and Odhav studied the antibacterial activity of maackiain (a flavonoid) and showed that it was active against *S. pneumoniae* with an MIC value of 12.5 μg/ml (Thiriloshani and Bharti, 2013).

Apart from those compounds with antibacterial properties, our results also unearthed a number of flavonoids and other phytochemicals with a variety of effects. For example, our GC MS and LC MS/MS results showed that rotenone (m/z=395.29) was present in hexane and methanol extracts. The flavonoid, rotenone is an important broad-spectrum insecticide, piscicide and pesticide (Hayes, 1991). Myricitrin has antipsychotic properties. Myricetin (m/z=319.1773) is a flavonoid with antioxidant, antiviral, antithrombotic, antidiabetic, anti-atherosclerotic and anti-inflammatory properties (Semwal et al., 2016). Myricetin can also act as a neuroprotectant. Another important flavonoid in bark extract is apiin, which shows anti-inflammatory activity that is mediated through inhibition of nitric oxide synthesis and inhibition of *i*NOS expression. Phenolic compound (p-coumaric acid glucoside) with antibacterial properties was also present in the bark extract, along with a number of important steroids possessing anti-inflammatory properties. Studies by Z. Lou et al showed that p-coumaric acid inhibited the growth of Gram-positive and Gram-negative bacteria with MICs values ranging from 10 to 80 mg/mL (Lou et al., 2012). The glycoside, syringin (m/z =373.1493) have been shown to possess antidiabetic properties (Sundaram Chinna Krishnan et al., 2014).

## 5. CONCLUSIONS

In summary, the results presented here demonstrate that methanol and hexane extract of *Xeroderris stuhlmannii* (Taub.) Mendonca & E.P. Sousa possesses antibacterial properties due to the presence of one or more phytochemicals likely acting synergistically to exert an effect on pathogens. The important compounds with antibacterial activity include ursolic acid, kaempferide, oleanolic aldehyde, p-coumaric acid glucoside and caffeine. Thus, *Xeroderris stuhlmannii* (Taub.) Mendonca & E.P. Sousa is a promising herbal medicinal plant for treatment of stomach disorders in rural communities where modern drugs are expensive. Future experiments aimed at separation and isolation of individual compounds with antibacterial, as well as toxicity assays are needed to examine the mechanisms of action of these compounds.

## 6. AUTHOR INFORMATION

### 6.1 Corresponding Author

Correspondence should be addressed to Freeborn Rwere, PhD., Department of Chemistry, School of Natural Sciences and Mathematics, Chinhoyi University of Technology, Off Chirundu Road, P. Bag 7724, Chinhoyi, Zimbabwe.

Current Address: Stanford University, School of Medicine, Department of Anesthesiology, Perioperative and Pain Medicine, 300 Pasteur Dr. Stanford, CA 94340, USA

### 6.2 Author Contributions

MAS and FR proposed the study and LB reviewed the research proposal. MAS and LK performed most of the experiments. FR and MAS performed data analysis and wrote the manuscript. All other authors contributed to experiments and manuscript writing. All authors have read and approved the final manuscript.

### 9. ACKNOWLEDGEMENT

The authors would like to acknowledge Prof. Grace Mugumbate, Chinhoyi University of Technology, for reading the manuscript and her valuable input in writing of the manuscript. The authors would also like to thank the Government Analyst Laboratories (GAL) in Harare, Zimbabwe and Mr. Daniel Zulu for GC MS analysis.

## REFERENCES

Aghakhani, F., Kharazian, N., Lori Gooini, Z., 2018. Flavonoid Constituents of Phlomis (Lamiaceae) Species Using Liquid Chromatography Mass Spectrometry. Phytochem. Anal. 29, 180–195. https://doi.org/10.1002/pca.2733

Asase, A., Oteng-Yeboah, A.A., Odamtten, G.T., Simmonds, M.S.J., 2005. Ethnobotanical study of some Ghanaian anti-malarial plants. J. Ethnopharmacol. 99, 273–279. https://doi.org/10.1016/j.jep.2005.02.020

Barbieri, R., Coppo, E., Marchese, A., Daglia, M., Sobarzo-Sánchez, E., Nabavi, S.F., Nabavi, S.M., 2017. Phytochemicals for human disease: An update on plant-derived compounds antibacterial activity. Microbiol. Res. 196, 44–68. https://doi.org/10.1016/j.micres.2016.12.003

Bonev, B., Hooper, J., Parisot, J., 2008. Principles of assessing bacterial susceptibility to antibiotics using the agar diffusion method. J. Antimicrob. Chemother. 61, 1295–1301. https://doi.org/10.1093/jac/dkn090

Cargnin, S.T., Gnoatto, S.B., 2017. Ursolic acid from apple pomace and traditional plants: A valuable triterpenoid with functional properties. Food Chem. 220, 477–489. https://doi.org/10.1016/j.foodchem.2016.10.029

Chen, Y., Yu, H., Wu, H., Pan, Y., Wang, K., Jin, Y., Zhang, C., 2015. Characterization and quantification by LC-MS/MS of the chemical components of the heating products of the flavonoids extract in Pollen typhae for transformation rule exploration. Molecules 20, 18352–18366. https://doi.org/10.3390/molecules201018352

Chithalen, J. V., Luu, L., Petkovich, M., Jones, G., 2002. HPLC-MS/MS analysis of the products generated from all-trans-retinoic acid using recombinant human CYP26A. J. Lipid Res. 43, 1133–1142. https://doi.org/10.1194/jlr.M100343-JLR200

Cock, I.E., Selesho, M.I., van Vuuren, S.F., 2019. A review of the traditional use of southern African medicinal plants for the treatment of malaria. J. Ethnopharmacol. 245, 112176. https://doi.org/10.1016/j.jep.2019.112176

Eumkeb, G., Siriwong, S., Phitaktim, S., Rojtinnakorn, N., Sakdarat, S., 2012. Synergistic activity and mode of action of flavonoids isolated from smaller galangal and amoxicillin combinations against amoxicillin-resistant Escherichia coli. J. Appl. Microbiol. 112, 55–64. https://doi.org/10.1111/j.1365-2672.2011.05190.x

Faraone, I., Rai, D.K., Chiummiento, L., Fernandez, E., Choudhary, A., Prinzo, F., Milella, L., 2018. Antioxidant activity and phytochemical characterization of senecio clivicolus wedd. Molecules 23, 1–17. https://doi.org/10.3390/molecules23102497

Fu, Z., Ling, Y., Li, Z., Chen, M., Sun, Z., Huang, C., 2014. HPLC-Q-TOF-MS/MS for analysis of major chemical constituents of Yinchen-Zhizi herb pair extract. Biomed. Chromatogr. 28, 475–485. https://doi.org/10.1002/bmc.3057

Grizzell, J.A., Echeverria, V., 2014. New Insights into the Mechanisms of Action of Cotinine and its Distinctive Effects from Nicotine. Neurochem. Res. 40, 2032–2046. https://doi.org/10.1007/s11064-014-1359-2

Haudecoeur, R., Peuchmaur, M., Pérès, B., Rome, M., Taïwe, G.S., Boumendjel, A., Boucherle, B., 2018. Traditional uses, phytochemistry and pharmacological properties of African Nauclea species: A review. J. Ethnopharmacol. 212, 106–136. https://doi.org/10.1016/j.jep.2017.10.011

Hayes, W.J., 1991. Handbook on Pesticides, Volume 1, 3rd ed. Academic Press.

Karygianni, L., Cecere, M., Argyropoulou, A., Hellwig, E., Skaltsounis, A.L., Wittmer, A., Tchorz, J.P., Al-Ahmad, A., 2019. Compounds from Olea europaea and Pistacia lentiscus inhibit oral microbial growth. BMC Complement. Altern. Med. 19, 1–10. https://doi.org/10.1186/s12906-019-2461-4

Keskes, H., Belhadj, S., Jlail, L., El Feki, A., Damak, M., Sayadi, S., Allouche, N., 2017. LC-MS-MS and GC-MS analyses of biologically active extracts and fractions from tunisian juniperus phoenice leaves. Pharm. Biol. 55, 88–95. https://doi.org/10.1080/13880209.2016.1230139

Lou, Z., Wang, H., Rao, S., Sun, J., Ma, C., Li, J., 2012. P-Coumaric acid kills bacteria through dual damage mechanisms. Food Control 25, 550–554. https://doi.org/10.1016/j.foodcont.2011.11.022

Maroyi, A., 2013. Traditional use of medicinal plants in south-central Zimbabwe_ review and perspectives _ Journal of Ethnobiology and Ethnomedicine _ Full Text.

Morais, L.C.S.L., Barbosa-Filho, J.M., Almeida, R.N., 1998. Central depressant effects of reticuline extracted from Ocotea duckei in rats and mice. J. Ethnopharmacol. 62, 57–61. https://doi.org/10.1016/S0378-8741(98)00044-0

Mugumbate, Grace C., Bishi, Lorraine Y., Rwere, F., 2018. Natural Products a Reservoir of Drugs for Treatment of Pulmonary Tuberculosis. EC Pulmonol. Respir. Med. 8, 545–553.

Navin, R., Mi Kim, S., 2016. Therapeutic Interventions Using Ursolic Acid for Cancer Treatment. Med. Chem. (Los. Angeles). 06, 339–344. https://doi.org/10.4172/2161-0444.1000367

Ngarivhume, T., Van’T Klooster, C.I.E.A., De Jong, J.T.V.M., Van Der Westhuizen, J.H., 2015. Medicinal plants used by traditional healers for the treatment of malaria in the Chipinge district in Zimbabwe. J. Ethnopharmacol. 159, 224–237. https://doi.org/10.1016/j.jep.2014.11.011

Njerua, S.N., Matasyoh, J., Mwaniki, C.G., Mwendia, C.M., Kobia, G.K., 2013. A Review of some Phytochemicals commonly found in Medicinal Plants. Int. J. Med. plants Cit. 105, 135–140.

Novotny, L., Abdel-Hamid, M.E., Hamza, H., Masterova, I., Grancai, D., 2003. Development of LC-MS method for determination of ursolic acid: Application to the analysis of ursolic acid in Staphylea holocarpa Hemsl. J. Pharm. Biomed. Anal. 31, 961–968. https://doi.org/10.1016/S0731-7085(02)00706-9

R., P., P., L., 2015. Qualitative screenings of photochemical and GC-MS analysis of ceropegia attenuata- an endangered tuberous plant. Int. J. Pharm. Sci. Res. 6, 2976–2981. https://doi.org/10.13040/IJPSR.0975-8232.6(7).2976-81

Semwal, D.K., Semwal, R.B., Combrinck, S., Viljoen, A., 2016. Myricetin: A dietary molecule with diverse biological activities. Nutrients 8, 1–31. https://doi.org/10.3390/nu8020090

Sledz, W., Los, E., Paczek, A., Rischka, J., Motyka, A., Zoledowska, S., Piosik, J., Lojkowska, E., 2015. Antibacterial activity of caffeine against plant pathogenic bacteria. Acta Biochim. Pol. 62, 605–612. https://doi.org/10.18388/abp.2015_1092

Sultana, B., Anwar, F., Ashraf, M., 2009. Effect of extraction solvent/technique on the antioxidant activity of selected medicinal plant extracts. Molecules 14, 2167–2180. https://doi.org/10.3390/molecules14062167

Sundaram Chinna Krishnan, S., Pillai Subramanian, I., Pillai Subramanian, S., 2014. Isolation, characterization of syringin, phenylpropanoid glycoside from Musa paradisiaca tepal extract and evaluation of its antidiabetic effect in streptozotocin-induced diabetic rats. Biomed. Prev. Nutr. 4, 105–111. https://doi.org/10.1016/j.bionut.2013.12.009

Thiriloshani, P., Bharti, O., 2013. Antimicrobial activity of plant phenols from Chlorophora excelsa and Virgilia oroboides. African J. Biotechnol. 12, 2254–2261. https://doi.org/10.5897/ajb12.961

